# Metformin protects the heart against chronic intermittent hypoxia through AMPK-dependent phosphorylation of HIF-1α

**DOI:** 10.1101/2024.01.11.575152

**Authors:** Sophie Moulin, Britanny Blachot-Minassian, Anita Kneppers, Amandine Thomas, Stéphanie Paradis, Laurent Bultot, Claire Arnaud, Jean-Louis Pépin, Luc Bertrand, Rémi Mounier, Elise Belaidi

## Abstract

Chronic intermittent hypoxia (IH), a major feature of obstructive sleep apnea syndrome (OSA), is associated with a more severe myocardial infarction. In this study, we performed RNA sequencing of cardiac samples from mice exposed to IH, which reveals a specific transcriptomic signature of the disease, relative to mitochondrial remodeling and cell death. Corresponding to its activation under chronic IH, we stabilized the Hypoxia Inducible Factor-1α (HIF-1α) in cardiac cells *in vitro,* and observed its association with an increased autophagic flux. In accordance, IH induced autophagy and mitophagy that is decreased in HIF-1α^+/_^ mice compared to wild-type animals suggesting that HIF-1 plays a significant role in IH-induced mitochondrial remodeling. Next, we showed that the AMPK metabolic sensor, typically activated by mitochondrial stress, is inhibited after 3 weeks of IH in hearts. Therefore, we assessed the effect of metformin, an anti-diabetic drug and potent activator of AMPK, on myocardial response to ischemia-reperfusion (I/R) injury. Daily administration of metformin significantly decreases infarct size without any systemic beneficial effect on insulin-resistance under IH conditions. The cardioprotective effect of metformin is lost in AMPKα2 knock-out mice demonstrating that AMPKα2 isoform promotes metformin-induced cardioprotection in mice exposed to IH. Mechanistically, we found that metformin inhibits IH-induced mitophagy in myocardium and decreases HIF-1α nuclear expression in mice subjected to IH. *In vitro* demonstrated that metformin induces HIF-1α phosphorylation, decreases its nuclear localization and subsequently HIF-1 transcriptional activity. Collectively, these results identify the AMPKα2 metabolic sensor as a novel modulator of HIF-1 activity. Our data suggest that metformin could be considered as a cardioprotective drug in OSA patients independently of their metabolic status.

## Introduction

Obstructive sleep apnea syndrome (OSA) is a highly prevalent chronic disease that affects nearly 1 billion people worldwide^1^. OSA is associated with the development or aggravation of cardiovascular and metabolic diseases^2^. Among cardiovascular diseases, OSA is linked to an increase in infarct size^3^ recognized to impair post-myocardial infarction (MI) structural and functional recovery^4^ and to accelerate the development of heart failure in apneic patients^5^.

OSA is characterized by repetitive pharyngeal collapses leading to chronic intermittent hypoxia (IH). IH is now well recognized to increase infarct size in several rodent models^6-8^. In myocardium, IH induces oxidative stress^8,9^ and sustained HIF-1 activation^6,7,,10^, leading to cardiomyocyte apoptosis through convergent mechanisms such as endoplasmic reticulum (ER) stress^7,11,,12^ and mitochondrial dysfunction^13,14^. Specifically, we previously demonstrated that chronic IH activates HIF-1, which is responsible for IH-induced ER stress^13-15^, calcium homeostasis alteration^7^, mitochondria associated ER-membrane disruption, mitochondrial dysfunction^13^ and cardiomyocyte death^6,7^.

Maintenance of mitochondrial function and integrity is primordial for cardiomyocyte survival, requiring a highly dynamic network that can alter metabolic function and the response to ischemia-reperfusion (I/R) injury^16^. Mitochondrial stress is linked to mitochondrial remodeling particularly through mitophagy ^17^. This process enables the selective clearance of dysfunctional mitochondria, correlating with beneficial effects for cardiac protection^18^. However, along severe stresses, mitophagy is inefficient, leading to the release of pro-apoptotic factors and cell death^19^. This illustrates that, depending on the context, short-term and long-term upregulation of mitophagy exert a differential impact on mitochondrial homeostasis and cell viability^20^.

Among the proteins activated independently by both mitochondrial stress and IH^21^, the AMP-activated Protein Kinase (AMPK) has been demonstrated to prevent cardiomyocyte death in several contexts^22^. However, the impact of chronic IH on myocardial AMPK activity remains unclear, depending on the depth and duration of IH, as well as the model and the tissue studied^23-26^.

Because chronic IH alters mitochondrial function and leads to cardiomyocytes apoptosis^13-15^, we first investigated if IH induced mitochondrial remodeling by evaluating autophagy and mitophagy. Since HIF-1 is recognized to activate genes encoding for the autophagic process^27,28^, we also questioned if IH-induced mitochondrial remodeling depends on HIF-1. Then, considering that AMPK activation has been demonstrated to be involved in myocardial response to mitochondrial stress, we questioned its myocardial activation state under chronic IH. We also investigated the impact of metformin treatment (an oral anti-diabetic drug and AMPK activator) on 1) myocardium response to ischemia-reperfusion (I/R) by measuring infarct size and 2) mitochondrial stress under IH. Finally, we explored how AMPK activation can modulate HIF-1 activity.

## Results

### Intermittent hypoxia exposure

Chronic IH was induced in mice as illustrating by the design in Fig.S1A. Chronic IH allowing 60 cycles (5% to 21% FiO_2_) *per* hour, is a unique and recognized model reproducing many cardiometabolic consequences of OSA. It was validated through systematic bodyweight assessments every week, as well as hematocrit measurement at the end of exposure. As expected, the bodyweight of hypoxic mice decreased during the first days of exposure (Fig.S1B) before rising normally, and hematocrit was significantly increased at the end of exposure (Fig.S1C).

### Intermittent hypoxia induces cardiac stress and mitochondrial remodeling

We investigated whether chronic IH alters the myocardial transcriptome (Fig. 1A). RNA sequencing and bioinformatic analysis revealed that 86 genes were dysregulated with 42 up-regulated and 44 down-regulated (FDR<0.05) (Fig.1B-C). Among the top 20 differentially expressed genes, Panther process analyses highlighted changes in erythrocytes O_2_/CO_2_ exchanges and oxidative stress, which are hallmark of IH (Fig.1D). Gene Ontology biological process analysis revealed changes in response to stress and metabolic processes (Fig.1E), consistent with the previously demonstrated impact of IH on myocardial mitochondrial dysfunction. Of particular importance, *Angpt1*, *Txnip*, *Nmrk2*, *Nuak1* or *Pfkfb1* genes belonging to the top 20 dysregulated genes are hallmark genes of hypoxia and glycolysis (Fig.1F). Finally, Gene Set Enrichment Analysis performed on the dysregulated genes revealed changes in cardiac stress, cell death and in mitochondrial remodeling biological processes (Fig.1G). Considering the pathways highlighted by our analysis and the impact of chronic IH on HIF-1-dependent mitochondrial dysfunction and cardiomyocyte death, we then investigated the impact of IH and HIF-1 on mitochondrial remodeling.

**Fig. 1:**
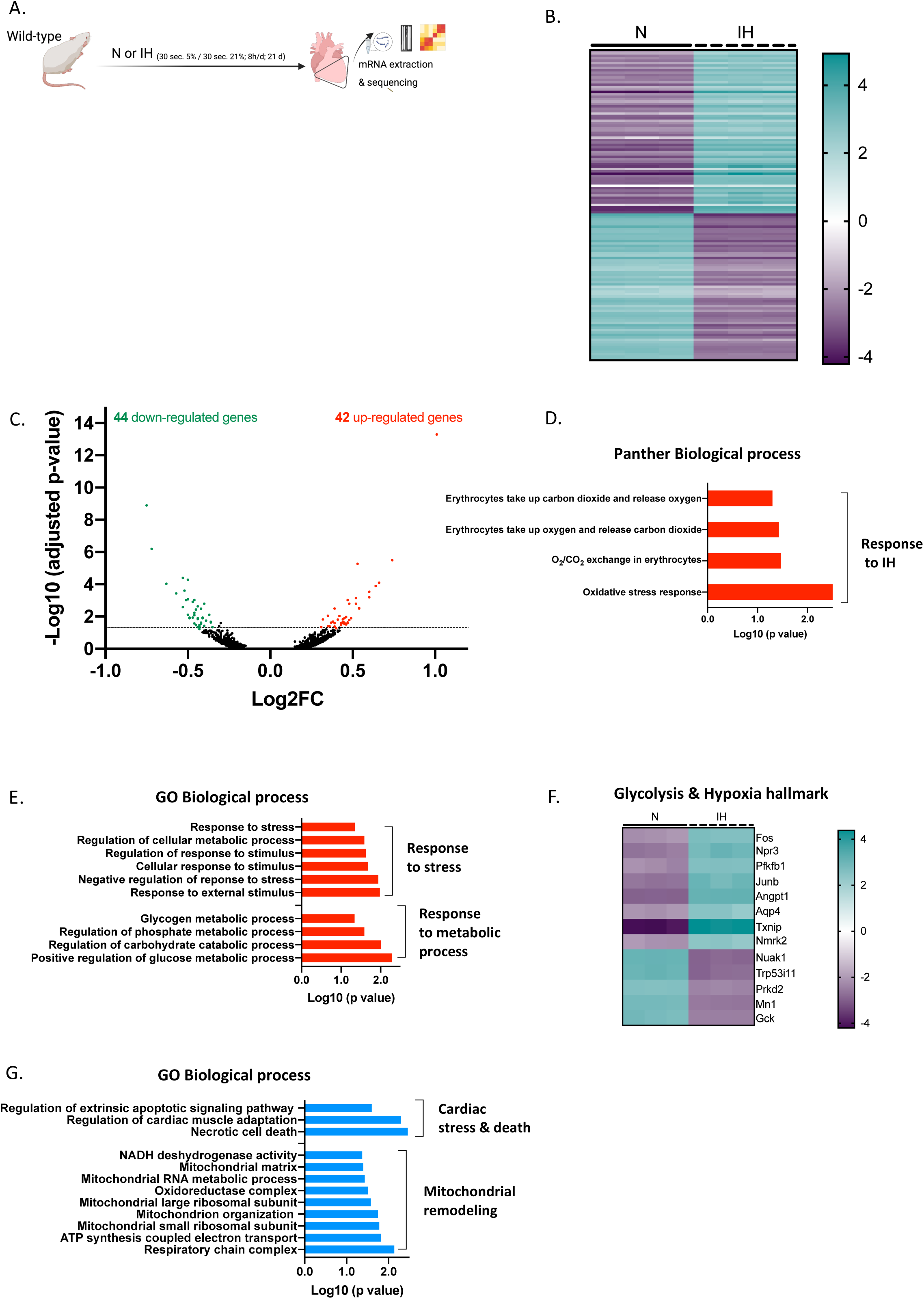
Intermittent Hypoxia unveils cardiac stress and mitochondrial remodeling in the heart through RNA-sequencing analysis. Flow chart of the RNA sequencing from mice exposed to Normoxia (N) or Intermittent Hypoxia (IH) (A). Heatmap showing differentially expressed genes (n = 3 biological replicates per group, FDR < 0.05) (B). Volcano plot of differentially expressed genes according to fold change and significance (FDR < 0.05) (C). (D-E) Enrichment analysis of biological process using Panther (D) and gene ontology (E) terms for top 20 differentially regulated genes. Heatmap showing glycolysis and hypoxia hallmark for top 20 differentially expressed genes (FDR < 0.05) (F). Enrichment analysis of biological process using gene ontology terms for all differentially expressed genes (G). Scheme were made with Biorender.com.

### Intermittent hypoxia-induced mitophagy is dependent on HIF-1

First, we investigated the link between HIF-1 chemical activation and autophagic flux in the *in vitro* H9c2 cardiomyoblast model (Fig.2A). HIF1**α** stabilization by cobalt chloride (CoCl_2_) increased HIF-1**α** nuclear localisation (Fig.2B-C) and this was associated with a marked increase in autophagic flux. Indeed, chloroquine strongly increased LC3II/I ratio in CoCl_2_-treated cells compared with control cells (Fig.2D).

**Fig. 2:**
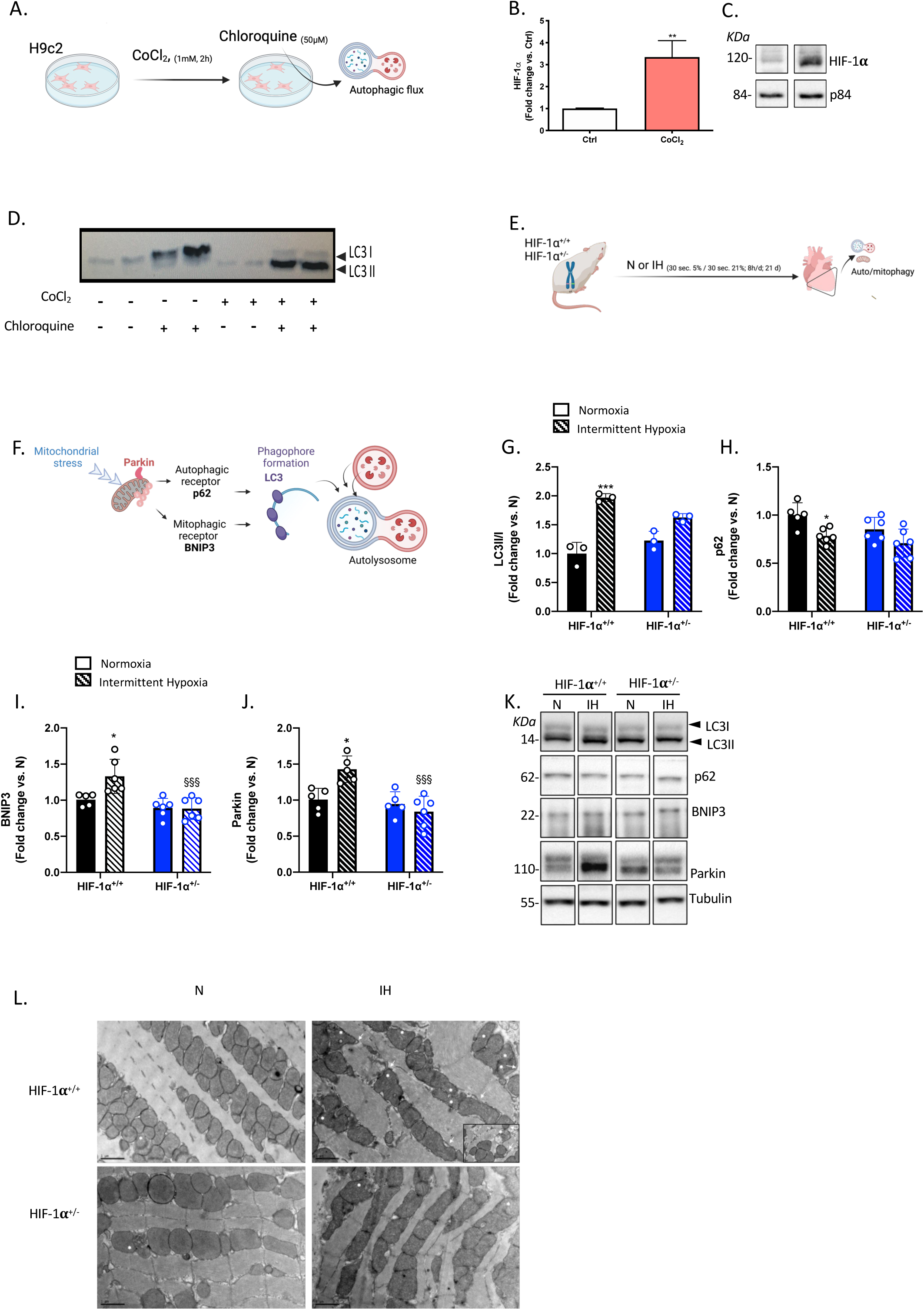
Intermittent hypoxia induces mitophagy through HIF-1 activation. (A-D) HIF-1 activation is associated with an increase in autophagic flux. H9c2 cells were treated with CoCl_2_ (1mM, 2h) with or without chloroquine (50 μM, 1h) (A), nuclear HIF-1**α** expression relative to p84 (B) and representative image of Western-blot (C), LC3 II and I Western-blots (D), *\*\*p<0.01, t-test*. **(E-L) Intermittent Hypoxia (IH)-induced autophagy and mitophagy is abolished in HIF-1α^+/-^ mice.** HIF-1**α**^+/+^ and HIF-1**α**^+/-^ mice were exposed to 21 days of Normoxia (N) or IH (1-min cycle of FiO_2_ 5%-21%) (E), scheme of protein explored and involved in mitophagy (F), LC3 II relative to LC3 I ratio (G), p62 (H), BNIP3 (I) and Parkin (J) expressions; representative image of Western-blot (K); representative images of cardiac mitochondrial remodeling acquired by transmission electron microscopy (stars for mitochondria with abnormal forms; arrows for autophagosomal membrane formation) (x2900). *\*p<0.05, ***p<0.001 IH vs N, ^§§§^p<0.001 HIF-1***α***^+/-^ vs HIF-1***α***^+/+^, two-way ANOVA, Šidák post-hoc tests.* Scheme were made with Biorender.com.

Next, to demonstrate the role of HIF-1 in IH-induced mitochondrial remodeling *in vivo*, we exposed HIF-1**α**^+/+^ and HIF-1**α**^+/-^ mice to normoxia or IH (Fig.2E). The partial decrease in HIF-1**α** mRNA transcripts (Fig.S2A) was sufficient to decrease HIF-1 activity in HIF-1**α**^+/-^ mice compared to HIF-1**α**^+/+^ upon IH (Fig.S2B). In HIF-1**α**^+/+^ and HIF-1**α**^+/-^ mice exposed to chronic IH, we investigated the expression of myocardial autophagic and mitophagic markers (Fig.2F). Twenty-one days of IH induced autophagy and mitophagy, as revealed by a significant increase in the LC3II/I ratio, a significant decrease in p62 (Fig.2G-H) as well as a significant increase in BNIP3 and Parkin (Fig.2I-J) protein expressions (Fig.2K) in HIF-1**α**^+/+^. In HIF-1**α**^+/-^, IH could not increase LC3II/I ratio and p62 expressions compared to normoxic mice (Fig.2G-H). Moreover, only under IH, BNIP3 and Parkin expressions were significantly decreased in HIF-1**α**^+/-^ mice compared to HIF-1**α**^+/+^ mice, suggesting that mitophagy under IH is driven by HIF-1 (Fig.2I-K). After chronic IH, there was no significant difference in Mfn2, OPA1 and Drp1 expressions between HIF-1**α**^+/+^ and HIF-1**α**^+/-^ (Fig.S2C-G). Images acquired by transmission electronic microscopy displayed characteristic mitophagy marks primarily in HIF-1**α**^+/+^ cardiomyocytes from mice exposed to IH, revealing an abundance of mitochondria with abnormal forms and the detection of autophagosomal membranes surrounding the mitochondria, (Fig. 2L). These results indicate that IH stresses mitochondria and induces mitophagy which depends on HIF-1.

### Metformin reverses IH-increased in infarct size through AMPK activation

IH was previously demonstrated to activate AMPK in *gastrocnemius* skeletal muscle of mice ^21^. Considering the mitochondrial stress induced by IH in hearts, we then investigated AMPK activation in hearts from IH mice. Surprisingly, we observed that AMPK phosphorylation on Thr172, its activation site, was not increased but is rather decreased in myocardium from IH mice (Fig.3A-C). Thus, we used metformin to activate AMPK during N or IH exposure to explore its impact on cardiomyocytes death after an *in vivo* ischemia-reperfusion procedure (Fig.3D). Metformin, at the concentration of 300 mg.kg^-1^, has been already shown to activate AMPK signaling in mouse heart ^29^. Twenty-four hours after the last administration, such treatment successfully increased the phosphorylation of ACC on Ser79, the *bona fide* substrate of AMPK, under IH (Fig.3A, 3E-G). Remarkably, despite equivalent areas at risk (AAR) in all groups (Fig.3H), IH significantly increased area at necrosis (AAN) which was significantly reduced by metformin treatment in IH mice only (Fig.3I-J). These results indicate that metformin specifically protects hypoxic hearts against ischemia-reperfusion. Surprisingly, while metformin improved systemic insulin sensitivity in normoxic mice, it failed to improve it in IH mice, indicating that the cardioprotective effect of metformin does not involve a systemic metabolic effect in IH conditions (Fig.S3A-B).

**Fig. 3:**
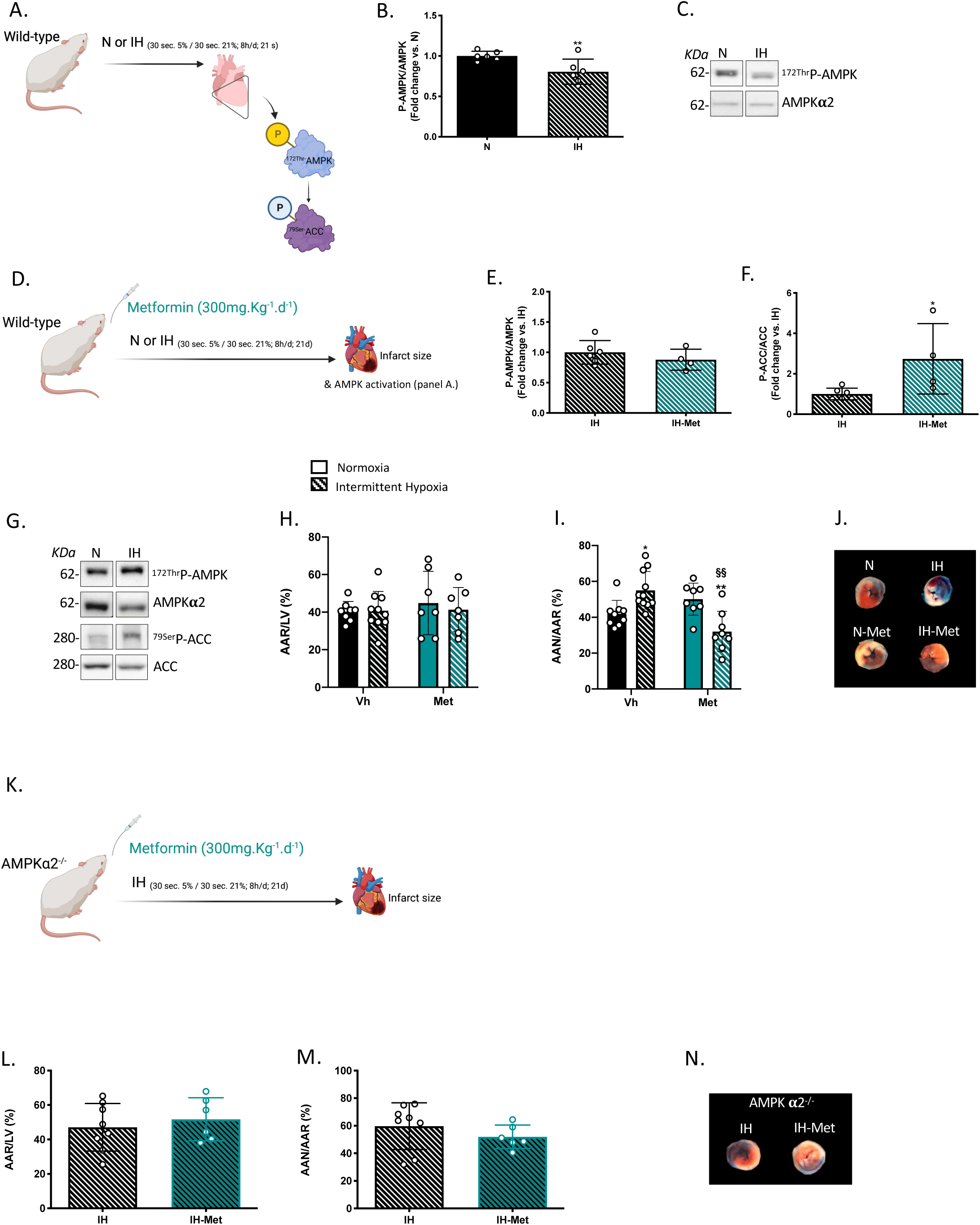
Metformin reverses intermittent hypoxia-induced increase in infarct size through AMPK activation. (A-C) **AMPK activation** (A) **is decreased in mice exposed to intermittent hypoxia (IH**). ^172Thr-^P-AMPK/AMPK (B) ratio and representative image of Western-blot (C), *\*\*p<0.01, t-test*. **(A, D-I) Metformin reverses IH-increase in infarct size through AMPKα2 activation.** Wild-type mice were treated with vehicle (Vh, CmCNa 0.01%, 0,1ml.10g^-1^) or metformin (Met, 300mg.kg^-1^.d^-1^) during exposure to 21 days of Normoxia (N) or IH (1-min cycle of FiO_2_ 5%-21%) (D); ^172Thr-^P-AMPK/AMPK (E)^79Ser-^PACC/ACC (F) ratio and representative image of Western-blot (G) in IH mice treated with metformin, *\*p<0.05, t-test;* mice were submitted to an ischemia-reperfusion protocol (D), area at risk (AAR) relative to Left ventricle (LV) (H) and area at necrosis (AN) relative to AAR (I) were calculated; representative colored slices (blue is the non-infarcted area, white is dead cells, red is viable cells) (J), *\*p<0.05 IH vs N, ^§§^p<0.05 Met vs Vh, Two-way ANOVA, Šidák post-hoc tests.* **(K-N) Metformin fails to protect the heart in AMPKα2^-/-^.** AMPK**α**2^-/-^ were exposed to 21 days of N or IH and were submitted to an ischemia-reperfusion protocol (K), AAR/LV (L), AAN/AAR (M) were calculated and representative slices (N), *Two-way ANOVA, Šidák post-hoc tests.* Scheme were made with Biorender.com.

Finally, we sought to determine whether the cardioprotective effect of metformin depends on AMPK activation by using AMPK**α**2^-/-^ (the main AMPK isoform in cardiomyocyte) mice exposed to IH and treated or not with metformin (Fig.3K). AMPK**α**2 expression is highly decreased in AMPK**α**2^-/-^ as compared to AMPK**α**2^+/+^ (Fig.S3C-D). Importantly, metformin failed to decrease infarct size in AMPK**α**2^-/-^ mice subjected to IH (Fig.3L-N). Taken together these data indicate that metformin rescues IH-induced increase in infarct size through AMPK**α**2 activation.

### Metformin reverses IH-induced mitophagy and decreases HIF-1 α nuclear expression

To comprehend how metformin specifically protects from IH-induced cardiomyocytes death, we investigated its effect on IH-induced auto/mitophagy (Fig.4A). Metformin significantly decreased LC3II/I ratio (Fig.4B), abolished IH-induced decrease in p62 expression (Fig.4C), and significantly decreased BNIP3 and Parkin expressions (Fig.4D-F) in the hearts of mice exposed to IH. Collectively, these results indicate that metformin reverses IH-induced mitochondrial stress. As done previously, we also checked the impact of metformin on other proteins involved in mitochondria dynamics (Fig.S4A-B). Except for an increase in OPA1 expression in both N and IH mice, metformin did not exert any effect on Mfn2 and Drp1 expressions (Fig.S4C-F). Because we demonstrated that IH-induced mitophagy is dependent on HIF-1, we also explored HIF-1**α** nuclear localization, a hallmark of its activation (Fig.4G). In hearts, IH significantly increased HIF-1**α** nuclear expression and this is significantly abolished by metformin (Fig.4H-I). Thus, our results suggest that the beneficial effect of AMPK activation by metformin can be explained by a decrease in HIF-1 activation.

**Fig. 4:**
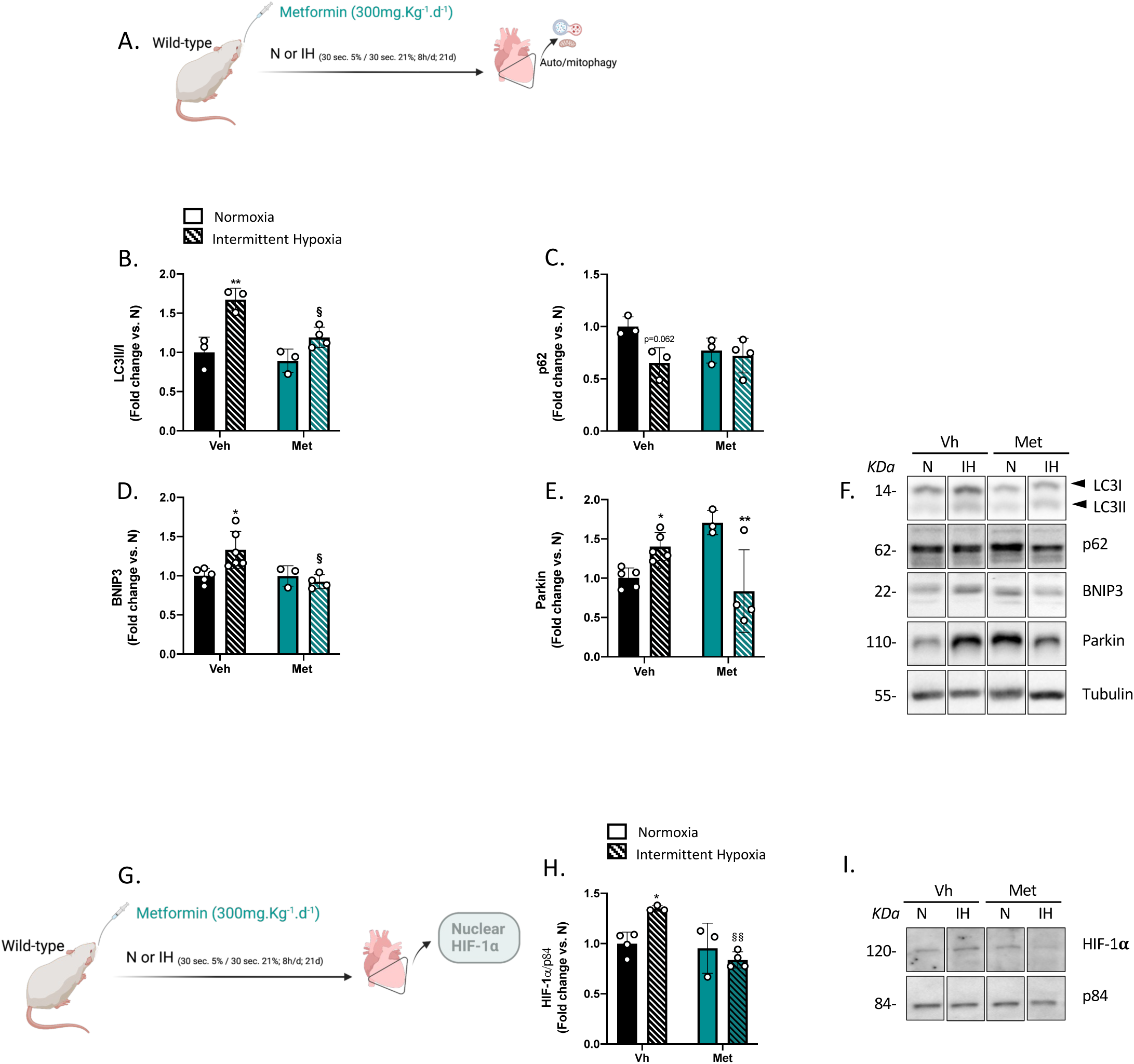
Metformin decreases intermittent hypoxia-induced mitophagy and HIF-1α nuclear localization. (A-F) Metformin decreases IH-induced mitophagy. Wild type mice were exposed to 21 days of Normoxia (N) or Intermittent Hypoxia (IH) (1-min cycle of FiO_2_ 5%-21%) and treated with vehicle (Vh, CmCNa 0.01%, 0,1ml.10g^-1^) or metformin (Met, 300mg.kg^-1^.d^-1^) (A), LC3 II relative to LC3 I ratio (B), p62 (C), BNIP3 (D), Parkin (E) expressions and representative images of Western-blot (F). **(G-I) Metformin decreases IH-induced increase in HIF-1α nuclear localization.** Nuclear HIF-1**α** expression was measured in N and IH mice treated or not with metformin (Met, 300mg.kg^-1^.d^-1^) (G), nuclear HIF-1**α** expression (H) and representative images (I). *\*p<0.05, **p<0.01 N vs IH, ^§^p<0.05, ^§§^p<0.01 Met vs Vh, two-way ANOVA, Šidák post-hoc tests.* Scheme were made with Biorender.com.

### Metformin increases HIF-1α phosphorylation and decreases HIF-1 activity

As HIF-1 phosphorylation is described to regulate its activity, we investigated the impact of metformin on HIF-1 phosphorylation in H9c2 cells exposed to CoCl_2_ (Fig.5A). As expected, CoCl_2_ increased HIF-1**α** expression and tended to increase HIF-1**α** phosphorylation (^Thr/Ser-P-^HIF-1**α**) in H9c2 control cells (Fig.5B-D). Interestingly, metformin significantly increased HIF-1**α** phosphorylation only in the presence of CoCl_2_ revealing that metformin-induced HIF-1**α** phosphorylation occurs when HIF-1**α** is stabilized (Fig.5C-D). In parallel, metformin significantly decreased HRE-GFP fluorescence in H9c2 which overexpress HIF-1**α**, indicating that metformin-induced HIF-1**α** phosphorylation could decrease HIF-1 activity (Fig. 5E-F). Finally, we treated AMPK**α**2^+/+^ and AMPK**α**2^-/-^ mice with CoCl_2_ in order to corroborate the role of AMPK**α**2 on HIF-1**α** phosphorylation (Fig.5G). CoCl_2_ tended to increase HIF-1**α** expression in AMPK**α**2^+/+^ and AMPK**α**2^-/-^ mice (Fig.5H). However, HIF-1**α** phosphorylation increased in AMPK**α**2^+/+^ but not in AMPK**α**2^-/-^ mice (Fig.5I-J), suggesting that AMPK**α**2 plays a role in HIF-1**α** phosphorylation. Taken together, these results indicate that AMPK**α**2 plays a role in HIF-1**α** phosphorylation and, that metformin can phosphorylate HIF-1**α** and decrease its activity.

**Fig. 5:**
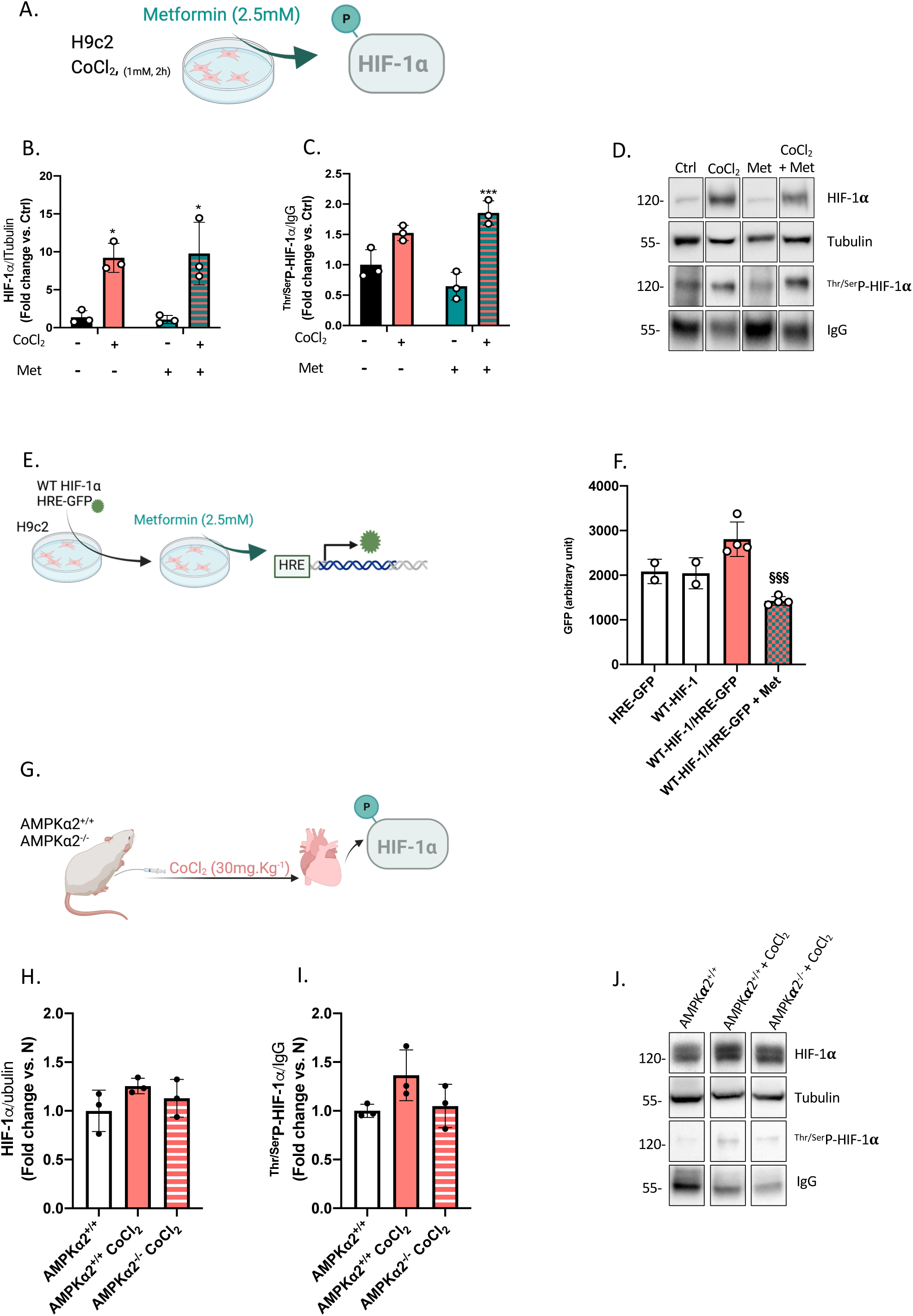
Metformin phosphorylates HIF-1α and decreases HIF-1 activity. **(A-D) Metformin induces HIF-1α phosphorylation.** H9c2 cells were treated with CoCl_2_ (1mM, 2h) and Vehicle (Vh, CmCNa, 0.1%) or metformin (Met, 2.5mM) (A), HIF-1**α** (B), ^Thr/ser-^P-HIF-1**α** (C) expressions assessed after immunoprecipitation and Western-blot; representative images (D) **(E-F) Metformin decreases HIF-1 activity.** H9c2 cells were co-transfected with WT-HIF-1 **α** and HRE-GFP plasmids and treated or not with Metformin (2.5mM) (E), fluorescence was measured using ⍰Ex at 475nm and ⍰Em spectrum 504-560nm (F), *\*p<0.05, ***p<0.001 CoCl_2_ vs Ctrl, two-way ANOVA, Šidák post-hoc tests; ^§§§^p<0.001 Met vs Vh*. **(G-J) AMPKα2 plays a role in HIF-1α phosphorylation.** AMPK**α**2^+/+^ and AMPK**α**2^-/-^ were treated with CoCl_2_ (30mg.kg^-1^) (G), HIF-1**α** (H), ^Thr/ser-^P-HIF-1**α** (I) expressions assessed after immunoprecipitation and Western-blot; representative image (J)*, one-way ANOVA, Tukey’s post-hoc tests.* Scheme were made with Biorender.com.

## Discussion

This study demonstrates that chronic IH induces an increase in infarct size through mitochondrial stress which is dependent on HIF-1 sustained activation. Our study also provides a rationale for a drug repositioning of metformin as a potential drug to specifically prevent cardiovascular risk in OSA.

Using an unbiased transcriptomic analysis, we uncovered that IH is associated with cardiac, oxidative and metabolic stress processes. Among the top 20 dysregulated genes by IH, HIF-1-related genes such as *Fos*, *Junb*, *Angpt1*, *Txnip*, *Pfkfb1* implicated in proliferation, oxidative stress or glycolysis were upregulated. Additionally, oxidative and metabolic stress were both contributors to cardiac stress and to mitochondrial remodeling, two biological processes highlighted in our non-biased approach.

We demonstrate that IH induces a mitochondrial stress as it activates autophagy and mitophagy, revealed by changes in auto/mitophagic markers, such as BNIP3 and Parkin. This is consistent with previous studies showing that chronic IH alters mitochondrial function and/or modify fuel usage in heart^13,14^. Furthermore, we showed that chronic IH-induced autophagy and mitophagy, is not observed in HIF-1**α** heterozygous mice demonstrating that IH-induced mitochondrial remodeling depends on HIF-1 upon IH. This aligns with the role of HIF-1 in inducing autophagy by activating the transcription of genes, including BNIP3^27,28^, one of the main autophagic receptor activated by hypoxia^30^. HIF-1 has also been shown to positively regulate parkin-mediated mitophagy in acute kidney injury^31^ and in diabetes^32^. In this study, we applied a chronic stress characterized by 21 days of 8 hours a day of IH. This stimulus is recognized to mimic severe OSA and induce pathologies associated with OSA^33,34^. Chronic IH is now well recognized to sustainably activate HIF-1^10^. In apneic patients at high cardiovascular risk, HIF-1**α** expression is positively correlated with the severity of OSA defined by the number of hypoxic cycles *per* hour^15^. Consistently, in pre-clinical models, HIF-1 has been demonstrated to be responsible for an increase in vascular remodeling^35^, mean arterial blood pressure, infarct size^6,7,,15^, cardiomyocyte apoptosis^15^, perivascular fibrosis and cardiac function alteration^36,37^. Indeed, HIF-1 has been shown to induce calcium homeostasis alterations^7,13^, ER stress ^7,38^ and more recently, to impair mitochondrial function^13,14^. All of these alterations, which are linked to HIF-1 upon IH, are potent stimulators of mitophagy, a dynamic process aiming to clear the mitochondrial pool^19^. This is consistent with the data provided by the transcriptomic analysis and the experiments performed in HIF-1**α**^+/-^ mice, demonstrating that HIF-1 is responsible for IH-induced myocardial mitochondrial stress that generates mitophagy.

Our current work reveals a decrease in AMPK signaling in the hearts of hypoxic mice. This result was unexpected considering that AMPK is a critical energy sensor activated by mitochondrial dysfunction, a decrease in ATP concentration and an increase in ADP-AMP/ATP ratio. Consistent with our result, a decrease in AMPK activation has been similarly reported in human adult cardiomyocytes exposed to chronic IH^39^. Inasmuch as the lack of explanation for such inhibition of AMPK by IH, we decided to investigate the effect of metformin, an AMPK activator widely used for efficiently treating type 2 diabetes^40^, that is also a pathology with a high incidence in apneic patients^41^. The cardiovascular benefits of metformin are a subject of controversy, depending on its efficiency in lowering glycemia^42^. Interestingly, we demonstrated that metformin significantly reduced infarct size in mice exposed to chronic IH without affecting systemic metabolism, as metformin treatment failed to improve systemic insulin resistance in IH mice. Importantly, under IH conditions, the cardioprotective effect of metformin is dependent on AMPK**α**2, as it failed to decrease infarct size in AMPK**α**2^-/-^ mice. These findings are of particularly significant, suggesting that metformin could be used exclusively as a cardioprotective agent in apneic patients. This highlights a potential repositioning of metformin for these patients, requiring a better understanding of the mechanisms involved in metformin cardioprotective effects under IH.

Maintaining energy homeostasis requires the coordination of a large array of compensatory mechanisms, some of which are triggered by AMPK activation. In rodents exposed to 6 to 10 weeks of IH, studies have demonstrated that AMPK activation by melatonin or adiponectin alleviates cardiomyocyte apoptosis and improves cardiac function by inhibiting oxidative stress^23^, inducing ER-phagy^26^ and activating autophagy and mitophagy^24,25^. However, these studies did not demonstrate any direct impact of AMPK activation on mitophagy. In those studies, AMPK activation indirectly induced mitophagy to protect against prolonged IH (6-10 weeks). These findings differ from our study in which mice were exposed to only 3 weeks of IH. In agreement with previous studies demonstrating a mitochondrial dysfunction in hearts from mice exposed to 3 weeks of IH^13^, we propose that mitochondria have already been submitted to a stress and have undergone remodeling characterized by an increase in mitophagy markers (without any modification in mitochondrial dynamics at this time). Interestingly, after 3 weeks, AMPK activation by metformin prevents the mitochondrial stress induced by IH, suggesting that earlier in the exposure (3 weeks vs 6-10 weeks), AMPK targets a key-pathway that is involved in the myocardial deleterious response to chronic IH. Thus, we hypothesized that AMPK activation could impact HIF-1 activity, involved in IH-induced mitochondrial remodeling after 3 weeks of IH. Furthermore, HIF-1 is the most important transcription factor activated by chronic IH^10,43,,44^, recognized to play a major role in myocardial susceptibility to ischemia-reperfusion^6,9,,13^ and mitochondrial dysfunction^13^.

Thus, we sought to determine if metformin can directly impact sustained HIF-1 activation induced by chronic IH. Here, we demonstrated that metformin abolished the IH-induced increase in HIF-1**α** nuclear localization, strongly suggesting that it can reduce its transcriptional activity^10,43^. HIF-1**α** primary sequence contains 15 threonine/serine phosphorylation sites playing a role in HIF-1**α** O_2_-dependent degradation, HIF-1**α** cytosolic retention^45^, HIF-1**α** dimerization and HIF-1 transcriptional activity^46^. We demonstrated *in vitro* that metformin significantly increases HIF-1**α** phosphorylation and decreases its activity when HIF-1 is sustainably stabilized by CoCl_2_ and overexpressed by transfection, respectively. Interestingly, HIF-1**α** phosphorylation failed to be increased by CoCl_2_ in AMPK**α**2 deficient mice, suggesting that among the proteins able to phosphorylate HIF-1**α**, AMPK**α**2 could participate. Some studies report that AMPK and HIF-1 decipher a reciprocal regulation in order to maintain cellular homeostasis^47^ but actually, very few studies indicate that HIF-1 activity can be modulated by AMPK and, only one work demonstrated a direct effect of AMPK on HIF-1 activity. Indeed, in *C. elegans*, Hwang et al. demonstrated that HIF-1**α** is phosphorylated by AMPK to reduce its activation^48^. Taken together, our results highlight that metformin blunts HIF-1 activity by increasing its phosphorylation, partially through AMPK**α**2 activation. This highlights the mechanism by which metformin could emerge as a drug, shielding the heart from the stress provoked by chronic IH.

In conclusion, our findings demonstrate that metformin efficiently counteracts the detrimental effects of chronic IH, providing a protection against cardiomyocyte death induced by ischemia-reperfusion. Beyond its established use for diabetes management, our data suggest that metformin may hold promise for patients with OSA and cardiovascular risks. Mechanistically, metformin triggers AMPK-dependent HIF-1**α** phosphorylation resulting in reduced HIF-1 activity and alleviation of mitochondrial stress (Fig. 6). This opens new avenues for exploring the potential effects of metformin as a modulator of HIF-1 activity in diverse pathologies including cancer and heart failure, where both AMPK and HIF-1 are involved.

**Fig. 6.**
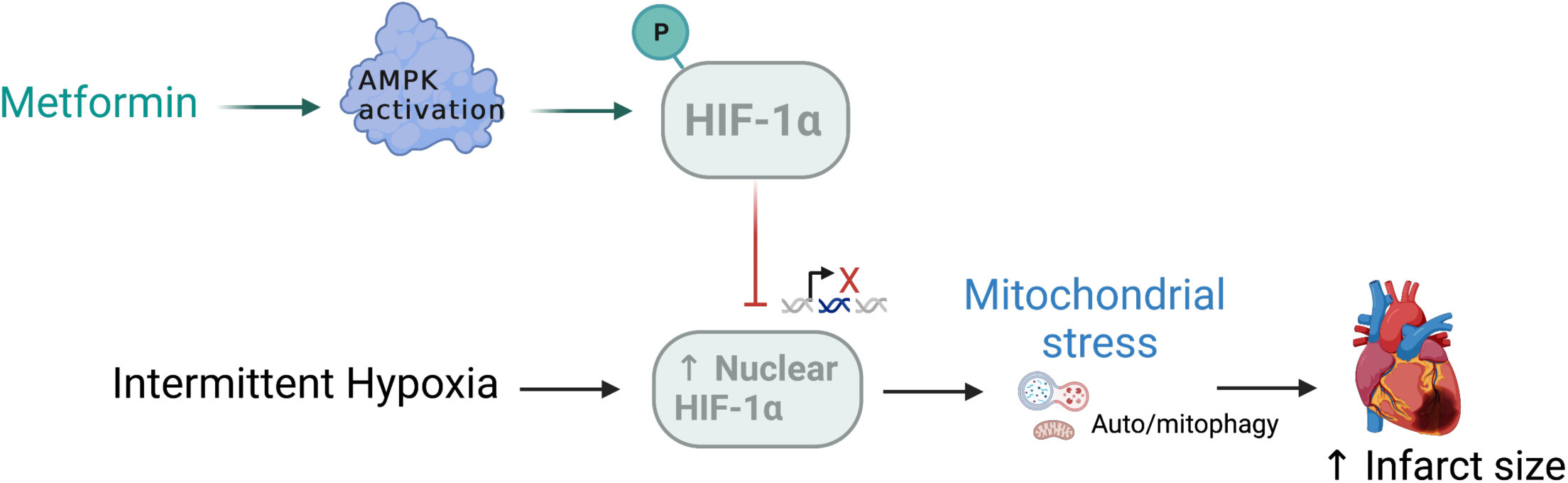
Graphical abstract Metformin reduces intermittent hypoxia (IH)-induced an increase in infarct size through AMPK α2. Metformin decreases HIF-1**α** nuclear localization, phosphorylates HIF-1**α** and decreases HIF-1 activity upon HIF-1**α** stabilization. The impact of AMPK**α**2 on HIF-1 activity could elucidate how metformin specifically protects hearts against ischemia-reperfusion injury, especially under chronic IH where, HIF-1 is known to induce mitochondrial stress and cell death. Scheme were made with Biorender.com

## Materials and Methods

### Materials

#### In vitro model, cell culture and design

H9C2 cardiomyoblasts were cultivated in classic conditions (DMEM high glucose, 10% SVF, 1% penicillin/streptomycin, 37°C, 5% CO_2_) in presence or absence of CoCl_2_ (1mM, 1h), used as a HIF-1α stabilizer. H9C2 were exposed to CoCl_2_ with or without chloroquine (50μM) in order to induce and assess autophagic flux. H9C2 were also exposed to CoCl_2_ in order to evaluate the effect of metformin treatment (2.5mM) on HIF-1α phosphorylation. Finally, the activity of HIF-1 was determined using an HRE-GFP plasmid transfected 24h before metformin treatment. This test was performed in control condition and in H9C2 simultaneously transfected with a WT-HIF-1α plasmid overexpressing HIF-1α. GFP-fluorescence intensity after metformin treatment was measured using Clariostar® fluorimeter (⍰Ex at 475nm and ⍰Em spectrum 504-560nm).

#### Rodent model of intermittent hypoxia

##### Animals

The protocol was approved by the French minister (APAFIS#23725-2020012111137561.v2). All experiments were performed in accordance with the European Convention for the Protection of Vertebrate Animals used for Experimental and Other Scientific Purposes (decree 2013-118 and orders of 1st February 2013); ARRIVE guidelines were followed. Male Swiss/SV129 mice with a heterozygous deletion of the gene encoding for the O_2_-sensitive HIF-1α subunit of HIF-1 (HIF-1α^+/-^) and their wild-type (HIF-1α^+/+^), aged 8-12 weeks were furnished by “Plateforme de Haute Technologie Animale”, Grenoble Alpes University. Male AMPKα2^-/-^ mice (C57/Bl6 background) were furnished by “Service Commun des Animaleries de Rockefeller” (Université Claude Bernard Lyon1). All animals were housed in the animal care facility of the HP2 laboratory (number: A38 516 10006). They were maintained on a 12/12 h light-dark cycle and had ad-libitum access to food and water.

##### Chronic intermittent hypoxia

After a week of acclimatation period, animals were exposed in their housing cages to 21 days of normoxia (N) or intermittent hypoxia (IH), 8 h per day during their sleeping period. The IH stimulus consisted in 60 s cycles alternating 30 s of hypoxia (hypoxic plateau at 5 % FiO2) and 30 s of N (normoxic plateau at 21 % FiO2). Normoxic mice were exposed to similar air-air cycles in order to avoid bias from noise and turbulences related to gas flow (Fig.S1A).

##### Experimental design

One week after arrival, mice were randomly assigned to be exposed to N or IH and 6 experimental sets were constituted. **In experimental set 1**, 2 groups were constituted (N=6 mice): HIF-1α^+/+^ -N, HIF-1α^+/+^ -IH (n=3 per group). Set 1 was used for RNA-seq analysis. **In experimental set 2**, 4 groups were constituted (N=24 mice): HIF-1α^+/+^ -N, HIF-1α^+/+^ -IH, HIF-1α^+/-^ -N and HIF-1α^+/-^ -IH. Set 2 was used for Western-blotting (n=3-6 per group) in order to assess mitochondrial stress and AMPK activation. **In experimental set 3**, 4 groups were constituted (N=48 mice): Vh-N, Vh-IH, Met-N and Met-IH; mice were treated daily with vehicle (Vh, CmCNA 0.1ml.10g^-1^.d^-1^) or metformin (Met, 300mg.kg^-1^.d^-1^) in order to perform *in vivo* ischemia-reperfusion (I/R) and to measure infarct size (n=12 per group). **In experimental set 5**, same groups than in experimental set 4 were constituted (N=24 mice) in order to perform an insulin tolerance test after 14 days of exposure and Western-blot after 21 days (n=3-6 per group). In **experimental set 6**, 2 groups were constituted (N=16 mice): IH and IH-Met; AMPKα2^-/-^ mice were exposed to IH and treated daily with Vh (CmCNA 0.1ml.10g^-1^.d^-1^) or Met (300mg.kg^-1^.d^-1^) in order to measure infarct size after I/R (n=8 per group). In **experimental set 7**, 3 groups were constituted (N=9 mice): AMPKα2^+/+^ and AMPKα2^-/-^ mice treated with CoCl_2_ (30 mg.kg^-1^) 1h before heart sampling in order to perform biochemical analyses (n=3 per group).

**Design and exclusion criteria**: concerning the biochemical experiments, depending on the experiment and based on our experience and on our practical feasibility, some designs were built with n=3 to 6. No exclusion was performed after the immunoblotting. Concerning the I/R protocol, animals dead before the end of the reperfusion were excluded.

### Methods

#### RNA sequencing, data processing and differential expression analysis

Heart tissues (>100mg) were sent to Active Motif and total RNA was extracted and qualified. For each sample, 1000ng of total RNA was then used in Illumina’s TruSeq Stranded mRNA Library kit (Cat# 20020594). Libraries were sequenced on Illumina NextSeq 550 as paired-end 42-nt reads. Sequence reads were analyzed with the STAR alignment – DESeq2 software pipeline described in the Data Explanation document of the manufacturer was performed. Finally, enrichment analyses were conducted using Gene Ontology and Panther databases.

#### Western-blot

Frozen heart biopsies were homogenized (Precellys® 24, Bertin Technologies, Montigny le Bretonneux, France) in lysis buffer as previously described and depending of the subfractionment^15^. Protein concentration was determined using Bradford assay. Then, proteins (30-100 µg) were separated by SDS-PAGE (8-12 % acrylamide) and transferred to polyvinylidene fluoride membranes. Membranes were blocked (5 % non-fat milk or BSA), then incubated overnight at 4°C with following primary antibodies: HIF-1α, LC3II, LC3I, p62, BNIP3, Parkin, Mfn2, OPA1, Drp1, ^172Thr^P-AMPK, AMPKa2, ^79ser^P-ACC, ACC1/2 (1:1000) and tubulin or p84 (1:2000). After that, appropriate horseradish peroxidase-conjugated anti-IgG (1:2500) were incubated for 1h at room temperature. Enhanced chemiluminescence was performed with the ECL Western-Blot detection kit (Bio-Rad, California, USA) and videoacquisition (ChemiDoc™ XRS+ System, Bio-Rad, California, USA). Density of protein bands was analyzed (Image J, NIH, USA) for each sample and expressed relative to corresponding tubulin expression (loading control) and normalized to the appropriate control group. We performed western-blot at least 2 times for each sample.

#### Immunoprecipitation and Western-blot

H9c2 were lysed as previously described (Moulin Antiox 22) and 500μg of proteins were incubated with 2μg of ^Ser/Thr-^phospho-protein antibody overnight at 4°C with rotation. The next day, samples were incubated for 2h with Pure Proteome protein G magnetic beads at 4°C and elution was performed using Laemmli buffer. Then, Western-blot was performed to detect HIF-1α.

### In vivo ischemia-reperfusion

At the end of the IH exposure, mice were anesthetized by an i.p. injection of pentobarbital (60 mg·kg^-1^). Animals were intubated with a tidal volume of 0.2 ml and a breathing rate of 160 per minute. Body temperature was maintained at 37 °C. Then, pericardectomy were performed and a 7-0 silk suture was passed around the left interventricular artery. After 45 min of ischemia, the ligature was removed and the myocardium was reperfused for 90 min, according to a previous study showing that 90 min is enough to induce stable necrosis areas^49^. At the end of reperfusion, the coronary artery was briefly re-occluded, and 1 ml Unisperse blue pigment was injected intravenously to delineate the area at risk (AAR). Then, hearts were excised and cut into five 1 mm-thick transverse slices. Each slice was incubated for 20 min in a 1% triphenyltetrazolium chloride solution at 37 °C to differentiate infarcted from viable myocardial areas^7^. Extent of AAR and area of necrosis (AN) was blindly quantified by planimetric analysis (Image J software) and corrected by the weight of each slice^50^.

### Insulin Tolerance Test

Whole-body insulin sensitivity was assessed by intraperitoneal insulin tolerance test. Injections of insulin (0.5 units.kg^-1^ total body weight) (NovoRapid; Novo Nordisk) were performed in conscious mice fasted for 6 h. Glycemia was measured by tail bleeding at different times (15, 30, 45, 60 and 90 minutes after insulin injection) using a glucometer (One Touch verio®).

### Statistical analysis

All analyses were done with GraphPad prism Software^®^. Two-way ANOVA following by Šidák post-hoc tests was performed. One-way ANOVA following by Tukey’s post-hoc test was performed when it was appropriate. T-test was performed when 2 groups have to be compared. Tests were performed after normality was accepted (Shapiro-Wilkinson test). Statistical significance was set for p values <0.05.

## Supporting information

Moulin, Blachot-Minassian et al. Supplemental figures

Moulin, Blachot-Minassian et al. Supplemental figures legendes

## Acknowledgement

We thank Lola Graizeau, Justine Dontaines, Iris Steuckardt, Sophie Bouyon, Guillaume Vial and Jonathan Gaucher for their technical support. We also thank Benoit Viollet and Bruno Guigas for their scientific advices.

We thank the “Fédération Française de Cardiologie” and “Agir pour les maladies chroniques » for their Financial support.

## Interest disclosure

Authors have no competing interests related to this study

## Notes

### Competing Interest Statement

The authors have declared no competing interest.

