## Supplementary figures and images for "Metformin protects the heart against chronic intermittent hypoxia through AMPK-dependent phosphorylation of HIF-1α"

### Moulin, Blachot-Minassian et al. Supplemental figures

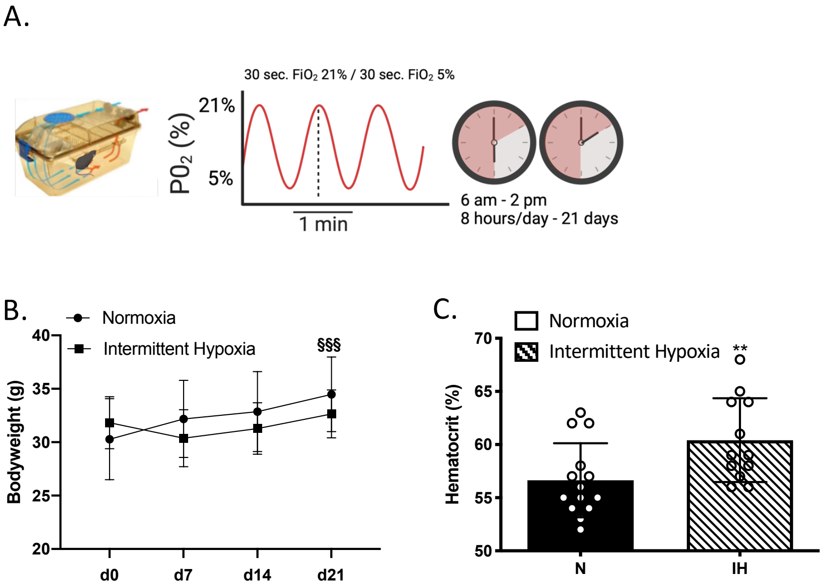

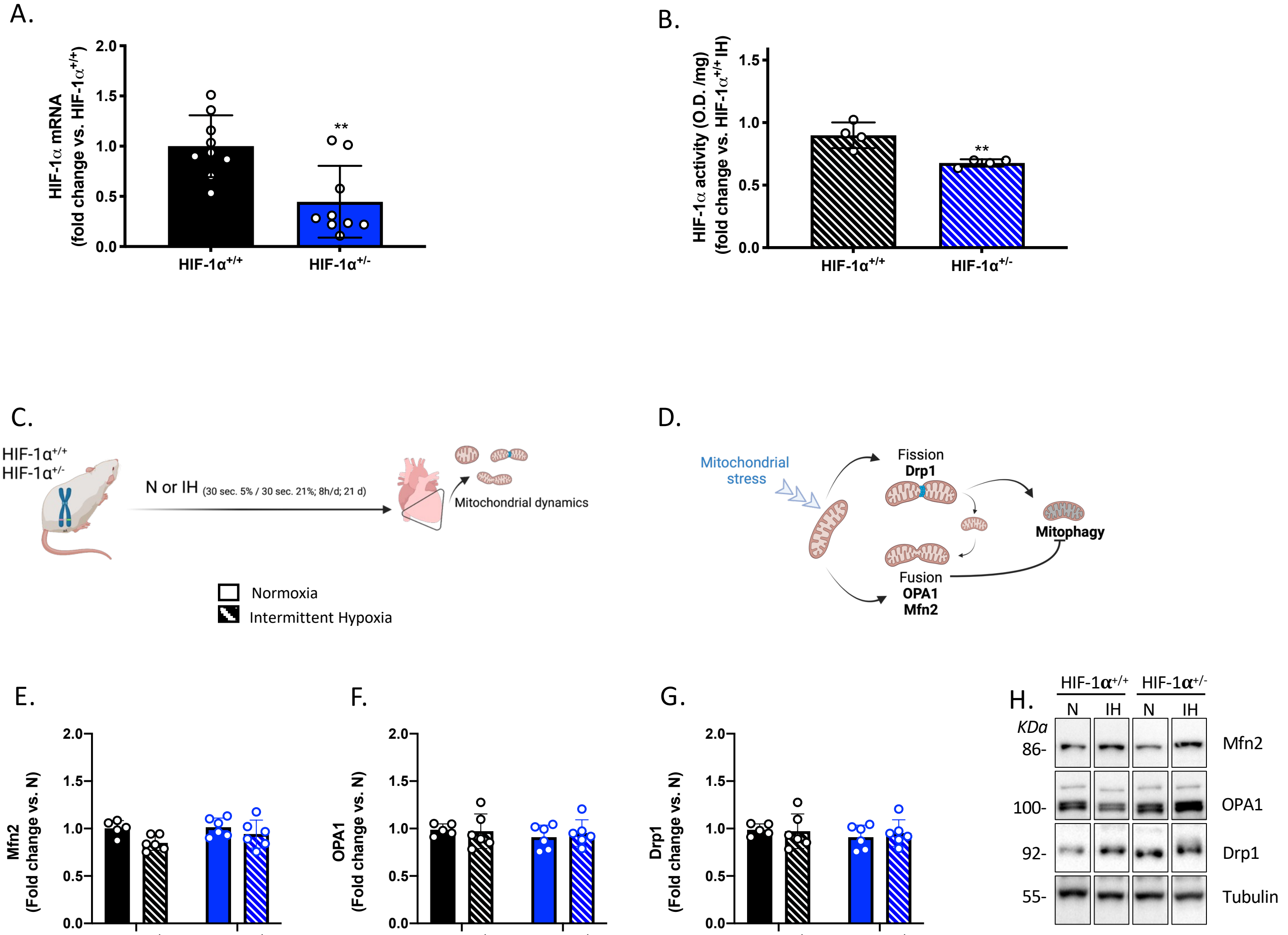

Figure S3

A.

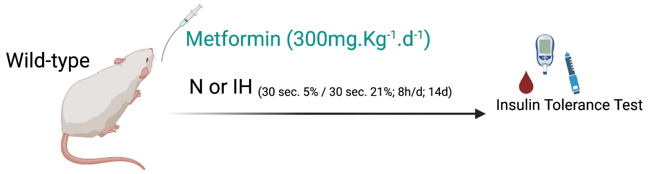


B.

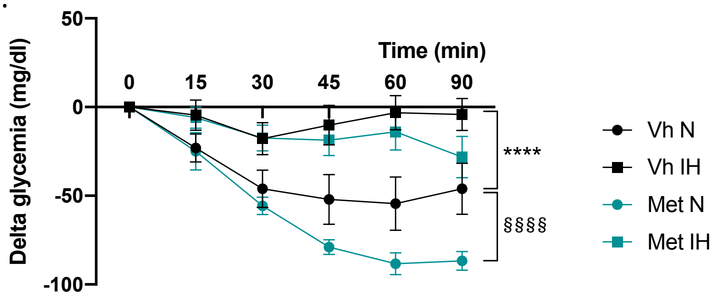


C.

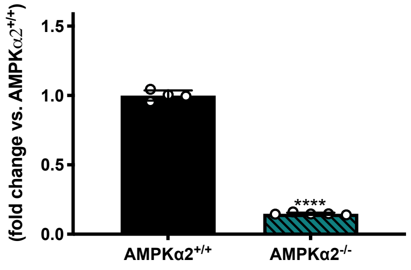


D.

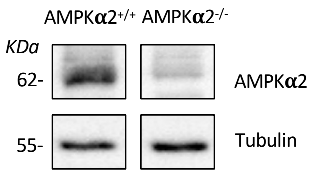

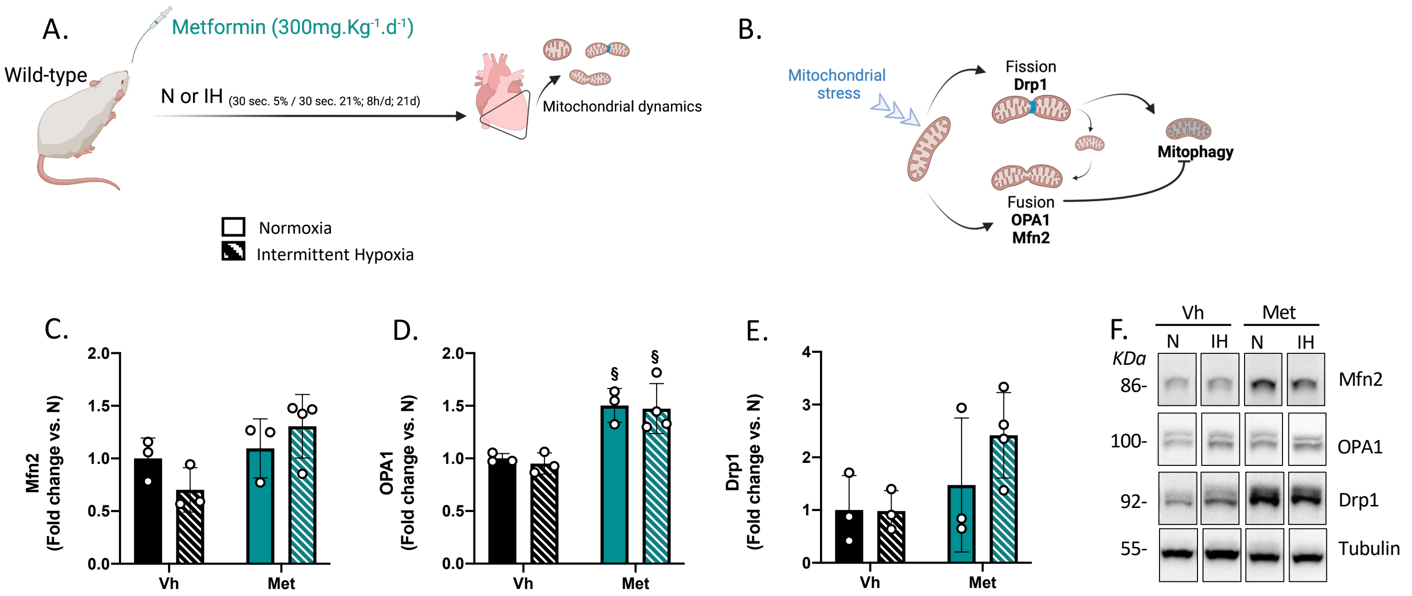
