## Supplementary material for "Metformin protects the heart against chronic intermittent hypoxia through AMPK-dependent phosphorylation of HIF-1α": Moulin, Blachot-Minassian et al. Supplemental figures legendes

**Supplemental figures legend:**

**Supplemental Fig. 1:**  **Intermittent hypoxia model validation**. Scheme of intermittent hypoxia (IH) exposition, animals were exposed in their own cages to 1-minute cycles of IH (30 sec. of inspired O_2_ fraction (FiO_2_) 21% / 30 sec. FiO_2_ 5%) or Normoxia (N, same noises generated by cycles without dropping FiO_2_), 8 hours per day (d), from 6 am to 2 pm (A). Bodyweight at d0, d7, d14 and d21 of N or IH (B). Hematocrit after 21 days of N or IH (C). *^§§§^p<0.001 d21 vs d0, two-way ANOVA,* Šidák *post-hoc tests; ^**^p<0.01 IH vs N, t-test.* Scheme were made with Biorender.com.

**Supplemental Fig. 2:** **(A-B) HIF-1𝛂 knock-out mice model validation.** HIF-1𝛂 mRNA expression in HIF-1𝛂^+/+^ and HIF-1𝛂^+/-^ (A). HIF-1 activity measured by ELISA (TRANS-AM Active Motif kit) in HIF-1𝛂^+/+^ and HIF-1𝛂^+/-^ exposed to IH (B). *^**^p<0.01 vs* HIF-1𝛂^+/+^*, t-test*. **(C-G) Mitochondrial dynamics in HIF-1𝛂^+/+^ and HIF-1𝛂^+/-^ mice** exposed to 21 days of Normoxia (N) or Intermittent Hypoxia (IH) (1-min cycle of FiO_2_ 5%-21%) (C), scheme of protein explored and involved in mitochondrial dynamics (D), Mfn2 (E), OPA (F), Drp1 (G), representative images of Western-blot (H). *Two-way ANOVA, Šidák post-hoc tests.* Scheme were made with Biorender.com.

**Supplemental Fig. 3: (A-B) Metformin fails to improve systemic insulin sensitivity in intermittent hypoxic (IH) mice.** Insulin tolerance test was performed with intraperitoneal injection of insulin (0.5mUI.kg^-1^) (A). Results are expressed in delta of glycemia from baseline before insulin administration (B). *^****^p<0.0001 IH vs N, ^§§§§^p<0.0001 Met vs Vh, Two-way ANOVA, Šidák post-hoc tests.* **(C-D) AMPK𝛂2 knock-out mice model validation**. AMPK𝛂2 expression in AMPK𝛂2 in AMPK𝛂2^-/-^ mice (C) and representative image of Western-blot (D). *^****^p<0.0001 vs* AMPK𝛂2 ^+/+^*, t-test*. Scheme were made with Biorender.com.

**Supplemental Fig. 4: Mitochondrial dynamics** in Wild-type mice exposed to 21 days of Normoxia (N) or Intermittent Hypoxia (IH) (1-min cycles of FiO_2_ 5%-21%) and treated or not with vehicle (Vh, CmCNa 0.01%, 0,1ml.10g^-1^) or metformin (Met, 300mg.kg^-1^.d^-1^) (A), scheme of protein explored and involved in mitochondrial dynamics (B), Mfn2 (C), OPA (D), Drp1 (E), representative images of Western-blot (F). *^§^p<0.05 Met vs Vh, two-way ANOVA, Šidák post-hoc tests.* Scheme were made with Biorender.com.
